# A Supervised Machine Learning Approach with Feature Selection for Sex-Specific Biomarker Prediction

**DOI:** 10.1101/2024.06.06.597741

**Authors:** Luke Meyer, Danielle Mulder, Joshua Wallace

## Abstract

Biomarkers play a crucial role in various aspects of healthcare, offering valuable insights into disease diagnosis, prognosis, and treatment selection. Recently, machine learning (ML) techniques have emerged as effective tools for uncovering novel biomarkers and improving predictive modelling capabilities. However, bias within ML algorithms, particularly regarding sex-based disparities, remains a concern. In this study, a supervised ML model was developed in order to predict 9 common biomarkers widely used in clinical settings. These biomarkers included triglycerides, body mass index, waist circumference, systolic blood pressure, blood glucose, uric acid, urinary albumin-to-creatinine ratio, high-density lipoproteins and albuminuria. During the validation test, it was observed that the ML models successfully predicted values within 5 and 10% error of the actual values. Out of the 121 female individuals tested, the following percentages of predicted values fell within this 10% range: 93% for albuminuria, 86% for waist circumference, 76% for BMI, and the lowest being 64% for systolic blood pressure and blood glucose. For the 119 male individuals tested, the percentages were as follows: 92% for albuminuria, 96% for waist circumference, 91% for BMI, 74% for blood glucose, and 68% for systolic blood pressure. Triglycerides, uric acid, urinary albumin-to-creatinine ratio and high-density lipoproteins all predicted lower than 50% for both male and female subgroups. Overall, the male subgroup had higher prediction scores than the female group.

**Graphical Abstract:** 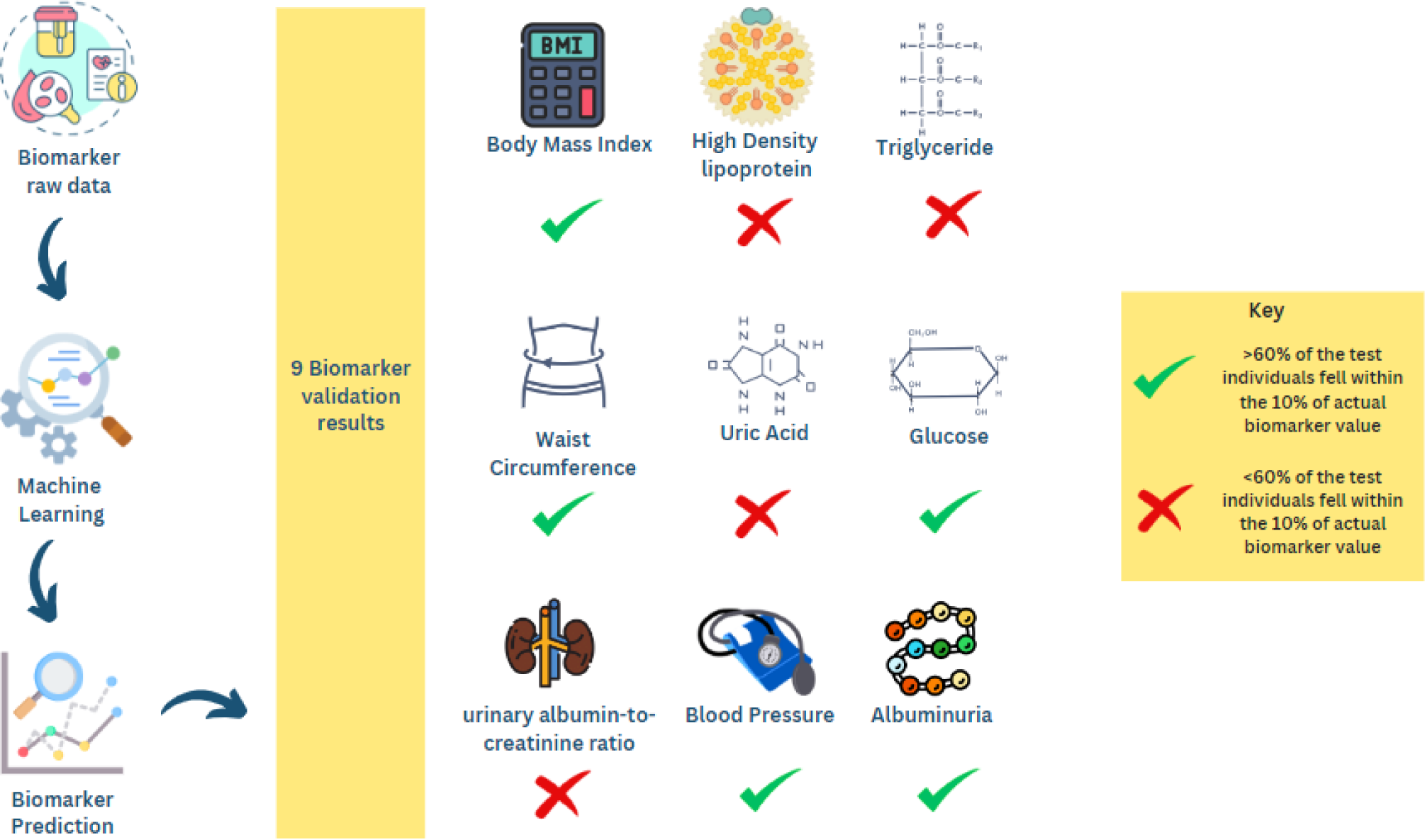

## 1. Introduction

The term “biomarker” refers to any medical signal that can be precisely measured and provides valuable insights into an individual’s medical state [1,2]. In healthcare, biomarkers are crucial for diverse aspects of patient care, playing pivotal roles in disease diagnosis, progression prediction, and treatment selection and monitoring. [3,4]. Due to the intricate interactions of physiological systems, biomarkers often exhibit correlations with each other, enabling more robust diagnostic and prognostic evaluations. Predictive biomarkers are particularly valuable as they facilitate the development of personalised therapeutic approaches and preemptive interventions [5]. The increasing focus on clinical trials driven by biomarker research aims at optimising disease management through personalised healthcare strategies such as screening and risk assessments. [4,6,7].

Recently, machine learning (ML) techniques have gained attention as effective techniques to uncover novel biomarkers [8–10]. ML algorithms are sensitive to high quality data and utilise a range of statistical, probabilistic, and optimisation techniques, drawing insights from prior knowledge to discern valuable patterns within vast, unorganised, and intricate datasets [11]. Big data also offers insight into relationships between various biomarkers and diseases, creating novel opportunities for predictive modelling in disease risk predictions [12]. By combining ML and extensive data sets one creates an opportunity for more accurate and reliable biomarker predictions.

In spite of the growing success of these ML methodologies, bias in biochemical algorithms is an overlooked issue, with one of them being sex based [13]. Straw and Wu, examined the ML predictions around liver disease and found that although the various studies and industries were able to predict liver disease in patients >70%, with optimisations done on ML models and features selected, they all failed to examine the effect that biological sex differences had on the ML prediction capability. Their study showed sex disparity in model performance for algorithms built from a commonly used liver disease dataset. They were also able to show how biochemical algorithms may reinforce and exacerbate existing healthcare inequalities [13]. Wang et al. conducted a ML study exploring the effect of sex bias in prognosis of lung cancers. Their conclusion was that more researchers in the cancer field are considering the effect of sex bias on cancers as females are found to have a better prognosis than males. They also state the necessity of developing sex-specific diagnosis and prognosis models [14]. Tokodi et al. considered mortality predictors among patients undergoing cardiac resynchronization therapy. Their in-depth analysis of features showed a marked sex difference in mortality predictors [15].

Taking into consideration that the exploration of biomarkers in healthcare, particularly through the lens of ML techniques, presents a promising avenue for revolutionising personalised medicine. It is imperative to address and mitigate biases within these algorithms, particularly concerning sex-based disparities, as highlighted by recent studies. Failure to account for such biases not only undermines the accuracy of predictive models but also perpetuates healthcare inequalities. Hence, the objective of this study was to develop accurate independent biomarker prediction models that account for sex-specific differences. To validate the accuracy of these predictions, standard ML evaluation techniques and diverse cross-validation statistical approaches were employed, thus affirming the strength and predictive capacity of the models.

## 2. Methods

### 2.1 Machine Learning Component

#### 2.1.1 Data Collection and Preprocessing

The study utilised SQL data derived from the National Health and Nutrition Examination Survey (NHANES), comprising 1931 participants, containing anthropometric and biomarkers associated with metabolic syndrome, type 2 diabetes (T2DM), and cardiovascular disease. Preprocessing techniques were implemented to ensure data accuracy and quality. Cases with missing information were systematically eliminated to prevent potential biases. Moreover, outliers surpassing 1.5 standard deviations from the average were removed to reduce distortions in subsequent analytical processes. Data normalisation was enabled in the hyperparameter tuning during the model setup procedure and individuals aged over 80 years old were excluded due to limited sample size. Following preprocessing, the cohort comprised 1199 participants, which was then utilised for training a series of supervised regression-based ML models. The study outline is indicated in Figure 1.

**Figure 1:**
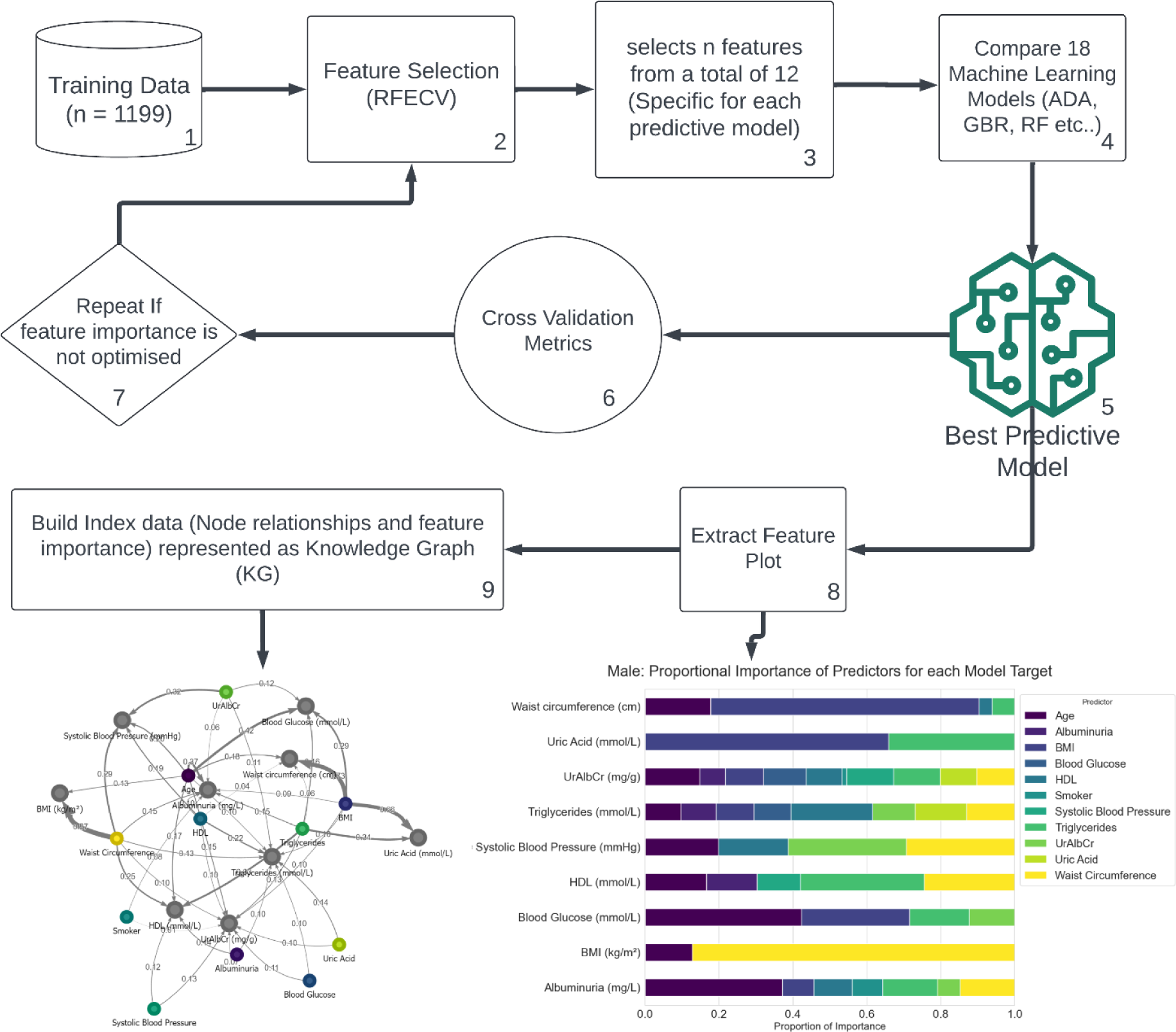
Summary of the biomarker prediction process

#### 2.1.2 Selection of Biomarkers (Clinical and Anthropometric)

For this study, we opted for a combination of anthropometric and clinical biomarkers to formulate a comprehensive predictive model applicable to both male and female subjects. In total, 14 features were initially considered for model training. These encompass five anthropometric indicators: sex, age, marital status, race, and smoking status; along with nine clinical biomarkers: blood glucose (mmol/L), HDL (mmol/L), triglycerides (mmol/L), and systolic blood pressure (mmHg), albuminuria (mg/L), UrAlbCr (mg/g), uric acid (mmol/L), waist circumference (cm), and BMI (kg/m²)

#### 2.1.3 Model Design and Training Procedure

We developed nine distinct biomarker predictive models using preprocessed data from NHANES. 80% of males were used for training and 20% were reserved for testing; the same method was applied to females in the cohort. In total, 483 females were allocated to the training set and 121 to the test set, while there were 476 males in the training set and 119 in the test set.

In our data preparation phase, we aggregated the dataset into male and female subsets to address sex-based disparities that could impact the predictive performance of our models, as evidenced by validation scores. These sex-related disparities sometimes led to instances of overfitting, where the model became overly tailored to specific sex patterns in the data. By isolating data by sex, we aimed to mitigate potential variations introduced by sex influences, consequently minimising overall model error.

During the training phase, we systematically compared various ML models (Table 1) using cross-validation metrics (as illustrated in Table 2) for each of the nine biomarker targets. During the model training process, optimised models are ranked for predictive accuracy and learning rate [16]. In our method, we used a diverse range of models including Linear, Ensemble, and Tree-Based models instead of relying on predetermined ones. This comprehensive approach enabled us to effectively capture intricate biological relationships that are difficult to capture with more traditional ML approaches.

**Table 1:**
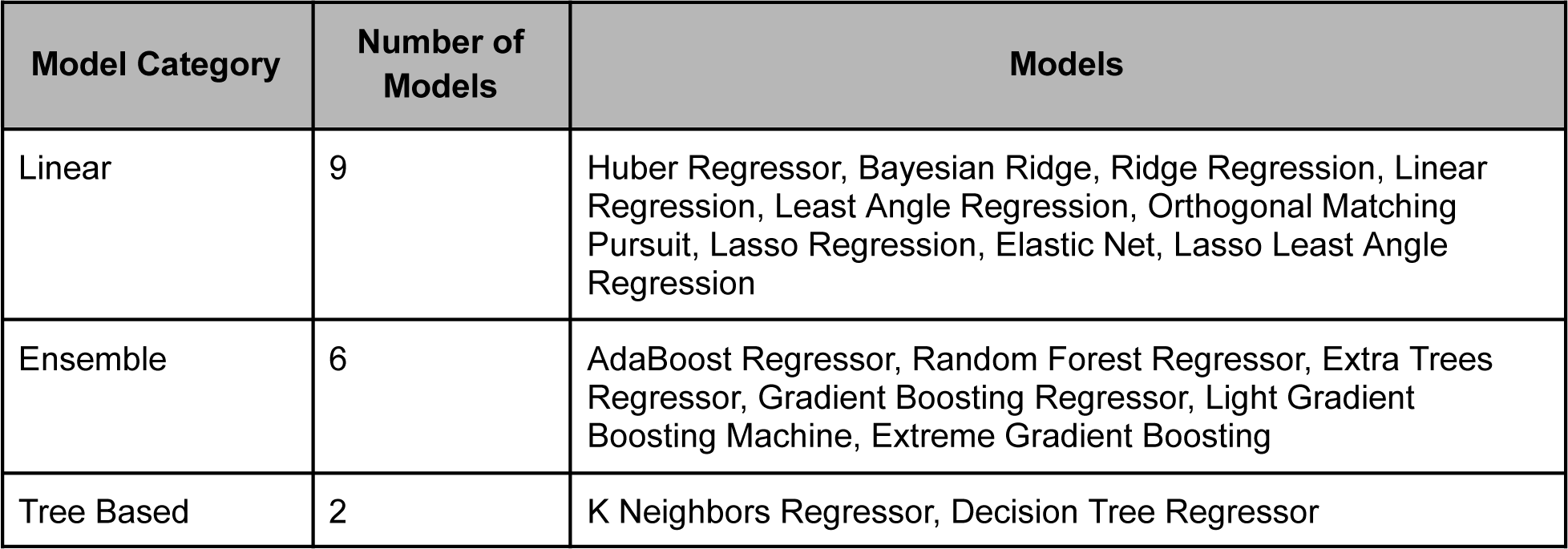
Showing the various model categories and models used in the study.

| Model Category | Number of Models | Models |
| --- | --- | --- |
| Linear | 9 | Huber Regressor, Bayesian Ridge, Ridge Regression, Linear Regression, Least Angle Regression, Orthogonal Matching Pursuit, Lasso Regression, Elastic Net, Lasso Least Angle Regression |
| Ensemble | 6 | AdaBoost Regressor, Random Forest Regressor, Extra Trees Regressor, Gradient Boosting Regressor, Light Gradient Boosting Machine, Extreme Gradient Boosting |
| Tree Based | 2 | K Neighbors Regressor, Decision Tree Regressor |

**Table 2:**
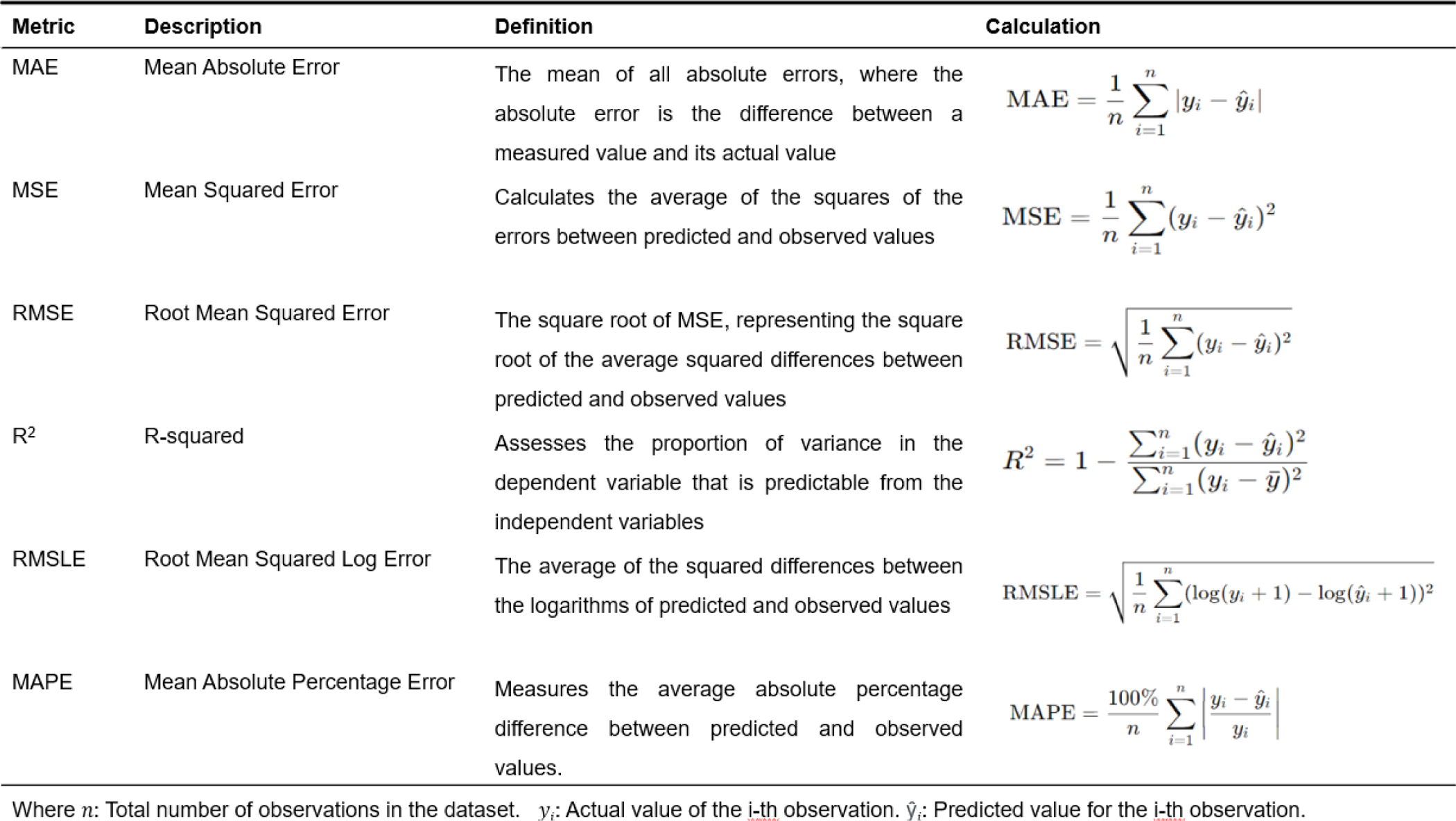
Cross validation metrics used in the study.

### 2.2 Optimisation of biomarker prediction - Feature selection

In order to enhance the effectiveness of the nine predictive models for the chosen biomarkers, a feature selection approach called recursive feature elimination with cross-validation (RFECV) was employed alongside supervised regression-based ML. [17]. This approach removes the need for predetermined feature selection during model training. The method involves iteratively removing less important features, training the model on remaining features, and then cross-validating to assess its performance.

### 2.3 Validation test on the hold-out set

A validation test was conducted on the 20% hold-out set. The aim of this test was to establish whether or not the predicted value fell within a ±5% or ±10% error margin of the actual value. The test was compiled visually as a heatmap which illustrates the counts of how many individuals in the hold-out set for males (119 individuals) and females (121 individuals) fell within the error margins. The percentage of these individuals was then calculated from these counts. The prediction was counted as acceptable if these percentages fell above 60% in the 10% error margin for an initial assessment.

## 3. Result

### 3.1 Clinical factors for the 1199 participants in the cohort

The descriptive statistics for the cohort were summarised in (Table 3). For each, the mean value was accompanied by the standard deviation in parentheses for continuous data and percentage for non-continuous data, providing an indication of the variability within the subgroup. The cohort exhibited a balanced sex distribution, encompassing diverse racial backgrounds and displaying homogeneity in age group representation. Notably, there was a higher prevalence of smokers within the male subgroup when compared to the female counterpart.

**Table 3:**
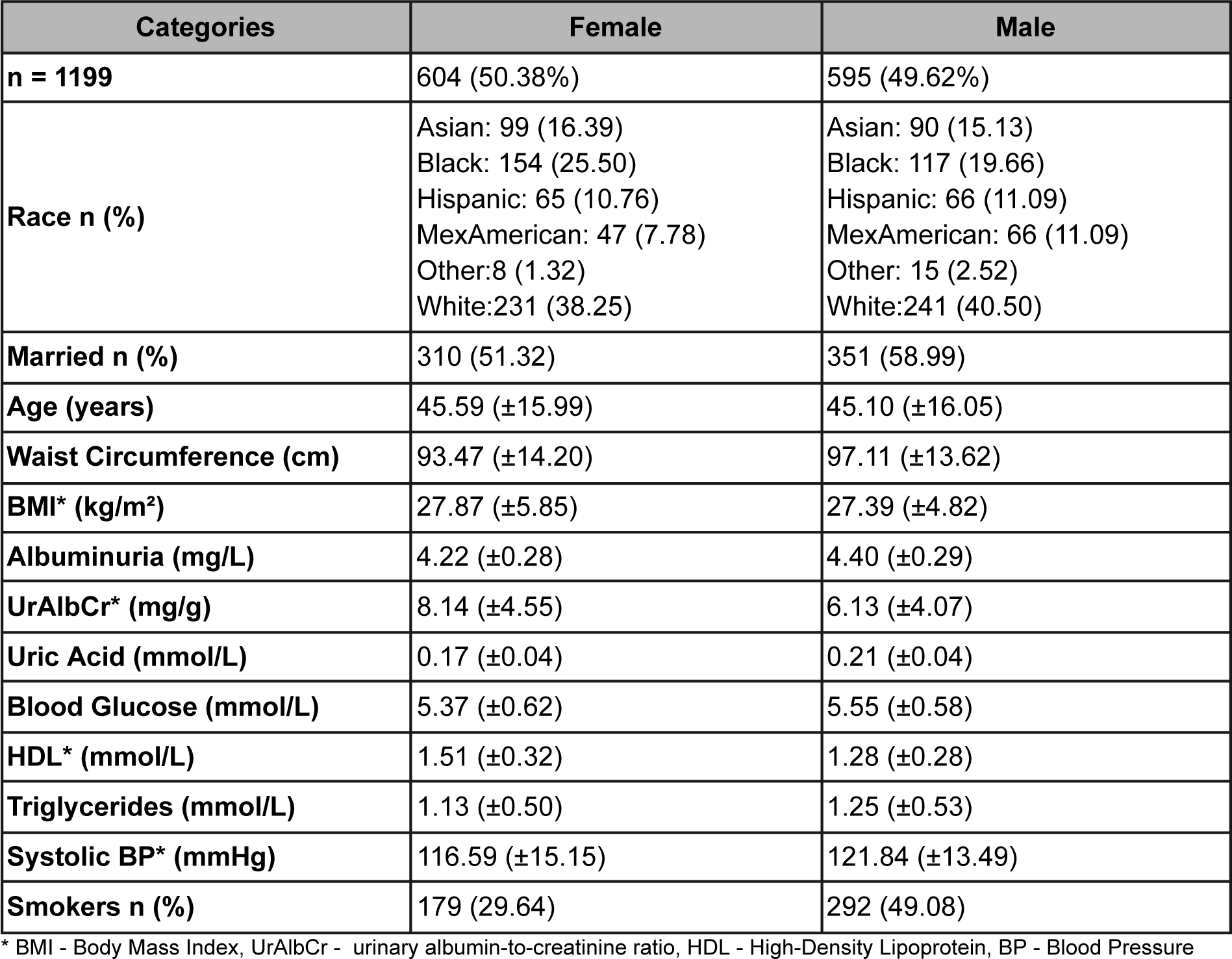
Clinical factors and demographic statistics and counts for the N=1199 participants in the NHANES cohort. Values indicated in brackets are the standard deviation for continuous data and percentage for non-continuous data.

| Categories | Female | Male |
| --- | --- | --- |
| **n = 1199** | 604 (50.38%) | 595 (49.62%) |
| **Race n (%)** | Asian: 99 (16.39)<br>Black: 154 (25.50)<br>Hispanic: 65 (10.76)<br>MexAmerican: 47 (7.78)<br>Other: 8 (1.32)<br>White: 231 (38.25) | Asian: 90 (15.13)<br>Black: 117 (19.66)<br>Hispanic: 66 (11.09)<br>MexAmerican: 66 (11.09)<br>Other: 15 (2.52)<br>White: 241 (40.50) |
| **Married n (%)** | 310 (51.32) | 351 (58.99) |
| **Age (years)** | 45.59 ($\pm 15.99$) | 45.10 ($\pm 16.05$) |
| **Waist Circumference (cm)** | 93.47 ($\pm 14.20$) | 97.11 ($\pm 13.62$) |
| **BMI\* (kg/m<sup>2</sup>)** | 27.87 ($\pm 5.85$) | 27.39 ($\pm 4.82$) |
| **Albuminuria (mg/L)** | 4.22 ($\pm 0.28$) | 4.40 ($\pm 0.29$) |
| **UrAlbCr\* (mg/g)** | 8.14 ($\pm 4.55$) | 6.13 ($\pm 4.07$) |
| **Uric Acid (mmol/L)** | 0.17 ($\pm 0.04$) | 0.21 ($\pm 0.04$) |
| **Blood Glucose (mmol/L)** | 5.37 ($\pm 0.62$) | 5.55 ($\pm 0.58$) |
| **HDL\* (mmol/L)** | 1.51 ($\pm 0.32$) | 1.28 ($\pm 0.28$) |
| **Triglycerides (mmol/L)** | 1.13 ($\pm 0.50$) | 1.25 ($\pm 0.53$) |
| **Systolic BP\* (mmHg)** | 116.59 ($\pm 15.15$) | 121.84 ($\pm 13.49$) |
| **Smokers n (%)** | 179 (29.64) | 292 (49.08) |

\* BMI - Body Mass Index, UrAlbCr - urinary albumin-to-creatinine ratio, HDL - High-Density Lipoprotein, BP - Blood Pressure

#### 3.1.2 Patterns in data variance

Analysis was done on the data to explore the variance in the data distribution for both male and female subgroups. Figure 2 shows the density distribution across the various biomarkers, whereas, Table 4 shows the statistical analysis done in order to determine the significant differences between mean and variances according to features between male and female groups.

**Figure 2:**
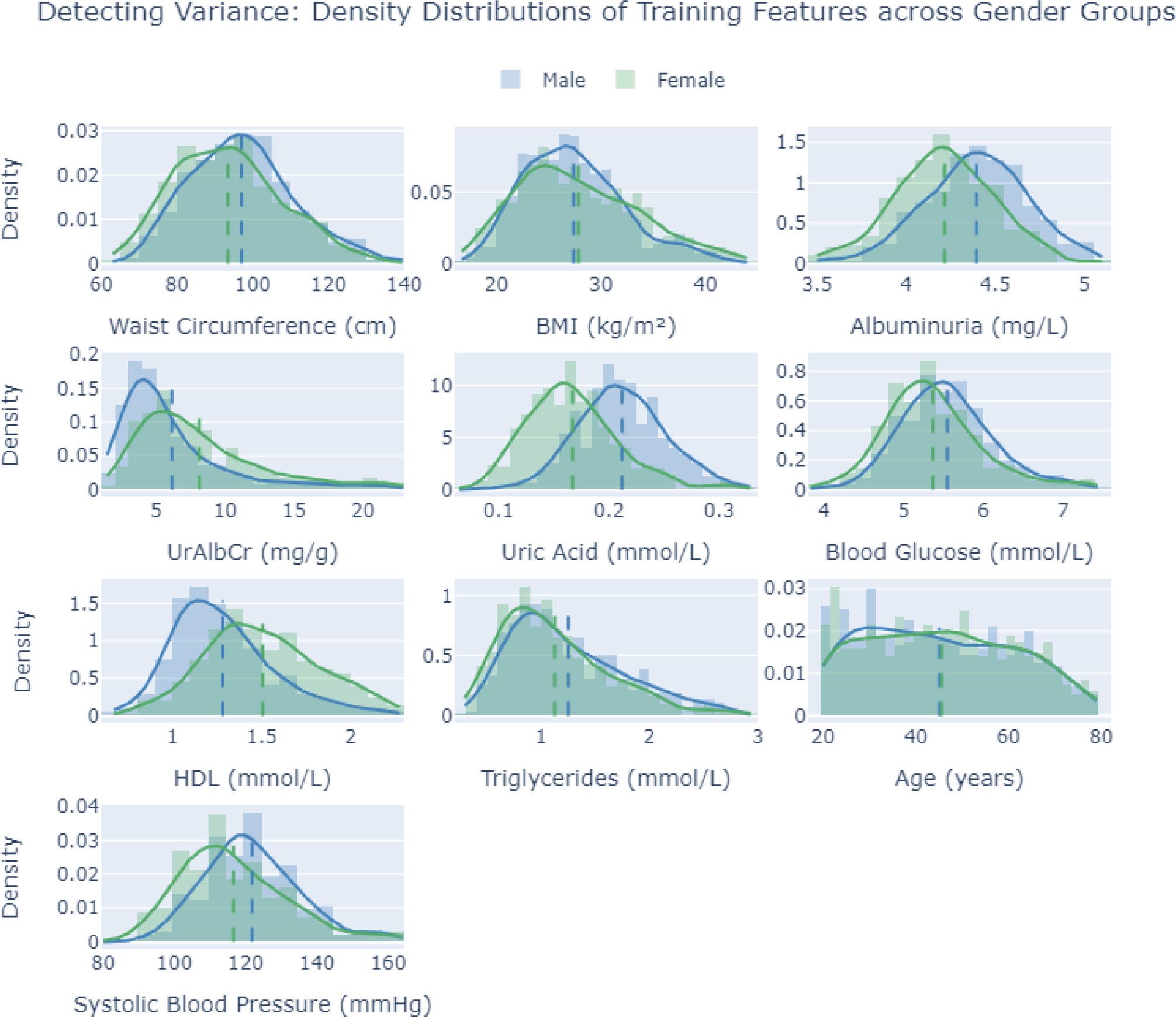
This figure displays the density distributions of various biomarkers across sex groups within the training dataset. Each subplot corresponds to a specific biomarker: Waist

**Table 4:** This table summarises the findings from a comparative analysis of various health biomarkers between male and female subgroups with respect to variance and mean values.

| Marker | *Levene's p-value ($\sigma^2$) | **t-test p-value |
| --- | --- | --- |
| Waist Circumference (cm) | 0.185 | p < 0.05 |
| BMI (kg/m<sup>2</sup>) | p < 0.05 | 0.121 |
| Albuminuria (mg/L) | 0.516 | p < 0.05 |
| UrAlbCr (mg/g) | p < 0.05 | p < 0.05 |
| Uric Acid (mmol/L) | 0.665 | p < 0.05 |
| Blood Glucose (mmol/L) | 0.436 | p < 0.05 |
| HDL (mmol/L) | p < 0.05 | p < 0.05 |
| Triglycerides (mmol/L) | 0.060 | p < 0.05 |
| Age (years) | 0.728 | 0.596 |
| Systolic Blood Pressure (mmHg) | p < 0.05 | p < 0.05 |

\*Levene's p-value which is designed to assess the equality of variances ( $\sigma^2$ ) between two groups. A p-value less than 0.05 indicates significant variance heterogeneity between male and female groups. \*\*t-tests are used to determine whether there are significant differences in the mean values of biomarkers between sexes. The type of t-test conducted (standard or Welch's) depends on the equality of variances assessed by Levene's test. \*\*\*A p-value less than 0.05 signifies a statistically significant difference in the mean, which is shown in red in the table..

Table 4. shows a variety of results indicated by Levene’s test (variance) and t-test (mean values), between male and female groups for the biomarkers shown in Figure 2. Importantly, these tests were not conducted on categorical data and therefore smoking status, race and marital status was excluded. The analysis was performed only on the continuous features used in the training data.

**Differences in both Variance and Means:** There was significant differences observed in both the variance (σ^2^) and mean values between the male and female subgroups for the systolic blood pressure (mmHg), HDL (mmol/L) and UrAlbCr (mg/g) biomarkers. Thus indicating that one subgroup had a wider range of values, which may imply greater diversity in how that biomarker was expressed or influenced by different factors within that sex.

**Differences in Means:** There was no significant differences in variance (σ^2^) between male and female groups for waist circumference (cm), albuminuria (mg/L), uric acid (mmol/L), blood glucose (mmol/L), and triglycerides (mmol/L). However, significant differences were seen for these markers in the mean values which was supported by the t-test. Thus indicating that the distribution of the biomarker values was consistent across genders, however, one gender tended to have higher or lower values for these biomarkers, which could highlight potential gender-specific health risks or conditions.

**Difference in Variance:** Levene’s test revealed significant variance differences in BMI (kg/m²) between male and female groups. The mean BMI (kg/m²) between the subgroups were not significantly different. This implies that this biomarker behaves similarly across genders, and gender may not be a critical factor for this measure. Although the average BMI is similar, the range of BMI values is different, indicating more variability in one gender.

### 3.2 Biomarker prediction optimisation including feature selection

RFECV was utilised on the complete set of 14 biomarkers to identify the optimal features for training individual models for each biomarker. This approach aimed to enhance prediction accuracy, minimise error, and optimise the number of features needed for an optimal validation score. Certain demographic factors such as race and marital status were omitted from the feature set due to their association with inaccuracies in the models. The study proceeded with analysis based on the remaining 12 features.

Table 3 presents the optimised sets of features for predicting various biomarker targets in both female and male subjects. For instance, the biomarker target “Albuminuria” was best predicted in females using body mass index (BMI), high-density lipoprotein (HDL), waist circumference, triglycerides, uric acid, and urine albumin-to-creatinine ratio (UrAlbCr). In contrast, the predictors for males included age, waist circumference, triglycerides, BMI, HDL, smoker status, and UrAlbCr. This pattern of varying optimal markers across sexes was observed across all listed biomarkers, illustrating the sex-specific differences in biomarker prediction.

Levene’s test results (Table 4) provided insights into sex-specific variability in biomarkers, which in turn influenced the selection of optimal features (Table 5). Data variance has successfully been shown to influence feature selection which could account for the differences in features seen in Table 5 [18–20].

**Table 5:** Features selected for biomarker optimisation.

| **Biomarker Target** | **Optimal Features for Females** | **Optimal Features for Males** |
| --- | --- | --- |
| **Albuminuria** | BMI, HDL, Waist circumference, Triglycerides, Uric acid, UrAlbCr | Age, Waist circumference, Triglycerides, BMI, HDL, Smoker, UrAlbCr |
| **Blood glucose** | Age, Waist circumference, Systolic blood pressure, HDL | Age, BMI, UrAlbCr, Triglycerides |
| **BMI** | Waist circumference, UrAlbCr, Age, Triglycerides, Albuminuria, Uric acid, Blood glucose, Systolic blood pressure, HDL, Smoker | Waist circumference, Age |
| **HDL** | Triglycerides, Age, Albuminuria, Waist Circumference, Blood glucose | Triglycerides, Waist circumference, Age, Albuminuria, Systolic blood pressure |
| **Systolic blood pressure** | Age, BMI, Blood glucose, UrAlbCr | UrAlbCr, Age, Waist circumference, HDL |
| **Triglycerides** | HDL, Waist circumference, Age, BMI, Uric acid, Albuminuria | Waist circumference, HDL, BMI, Albuminuria, Uric acid, Blood glucose, UrAlbCr, Age |
| **UrAlbCr** | Systolic blood pressure, Albuminuria, Waist circumference | Age, Systolic blood pressure, Triglycerides, Blood glucose, BMI, Waist circumference, Uric acid, HDL, Albuminuria, Smoker |
| **Uric acid** | BMI, Triglycerides, Albuminuria, Age, Waist circumference, Blood glucose | BMI, Triglycerides |
| **Waist circumference** | BMI, Age, Smoker, Triglycerides, Blood glucose, HDL, Albuminuria, Systolic blood pressure | BMI, Age, Triglycerides, HDL |

\*BMI - Body Mass Index, UrAlbCr - urinary albumin-to-creatinine ratio, HDL - High-Density Lipoprotein

Figure 3 serves as a complement to Table 5. Figures 3a and 3b depict feature plots for females and males, respectively, showcasing the weighted contributions of selected features to optimise specific biomarkers. Figures 3c and 3d present knowledge graphs for females and males, respectively, offering an alternative visual representation of how features contributed to various biomarkers, with heavier lines indicating higher feature contribution. Finally, Figure 3e provides a comprehensive overview of the actual counts of feature selections contributing to all nine biomarkers within separate female and male groups, as well as in the combined population.

**Figure 3:**
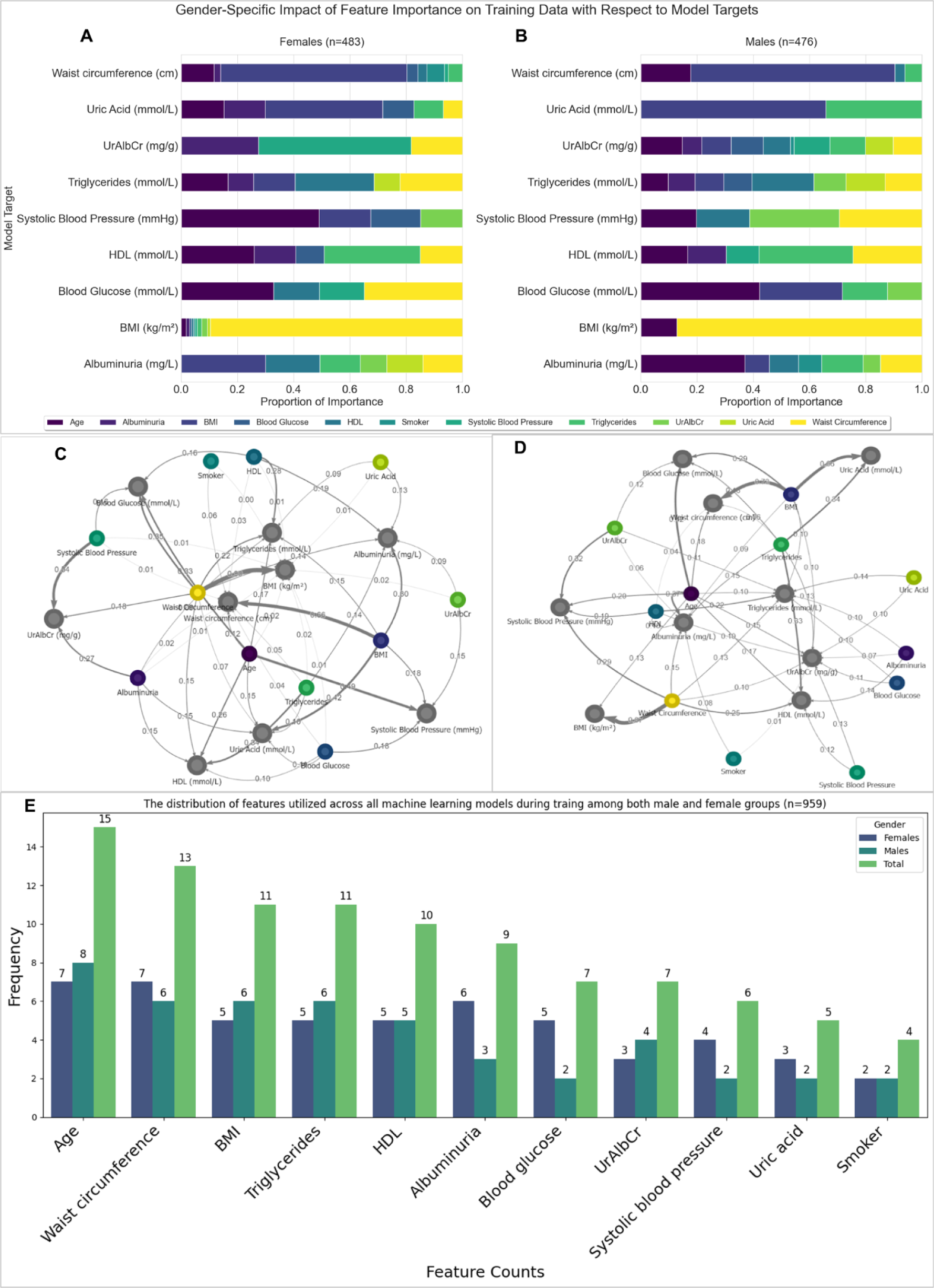
Showing the contribution of features for biomarker optimisation in the female and male subgroups. a, b) Proportional importance/weighting of features for biomarker optimisation in females and males respectively. c, d) Knowledge graphs visualising the features selected for biomarker optimisation in female and male subgroups. The coloured circles are the features, whereas the grey circles are the target biomarkers e) Frequency count for each biomarker that was included as a feature in the various biomarkers optimisation for males, females and combined subgroups.

In Figure 3a,c for the female subgroup, it can be seen that waist circumference was the highest contributing feature (weighting) for BMI biomarker prediction, age for systolic blood pressure and blood glucose, HDL for triglycerides, and triglycerides for HDL, systolic blood pressure for UrAlbCr, and BMI was the highest contributor to 3 of the biomarkers, namely albuminuria, uric acid and waist circumference. A different pattern was observed in the male subgroup (Figure 3b,d) where waist circumference contributed towards BMI and triglycerides, age for blood glucose, albuminuria and UrAlbCr, BMI for uric acid and waist circumference, UrAlbCr for systolic blood pressure, and triglycerides for HDL.

The knowledge graphs in Figure 3c,d assisted the feature map (Figure 3a,b) by showing the clear influence of the features to their biomarker targets. Considering Figure 3c, BMI had a larger contribution towards waist circumference, followed by uric acid, albuminuria, systolic blood pressure and lastly triglycerides. This trend was seen by the thickness of the arrows coming from the BMI feature. Similarly with age; age contributed more towards blood pressure, then followed by blood glucose, HDL, triglycerides, uric acid and finally waist circumference. In Figure 3d, waist circumference had a larger contribution towards BMI, followed by systolic blood pressure, HDL, albuminuria, triglycerides and finally UrAlbCr in the male subgroup. Age contributed more towards blood glucose and then albuminuria, HDL, waist circumference, BMI, triglycerides, and UrAlbCr.

In Figure 3e, age was the most frequently selected feature for both male and female groups in the ML models. For blood glucose, the analysis showed a higher frequency count in males compared to females. The analysis of feature selection across sex revealed distinct trends in feature frequency: In males, features related to adiposity measures (waist circumference, BMI, triglycerides) had higher frequency counts. In females, waist circumference and albuminuria frequently clustered together, while BMI, triglycerides, blood glucose, and HDL also tended to cluster.

The results in Table 6, showed the best ML model and corresponding metrics for the predictions of the various biomarkers for the two subgroups.

**Table 6:**
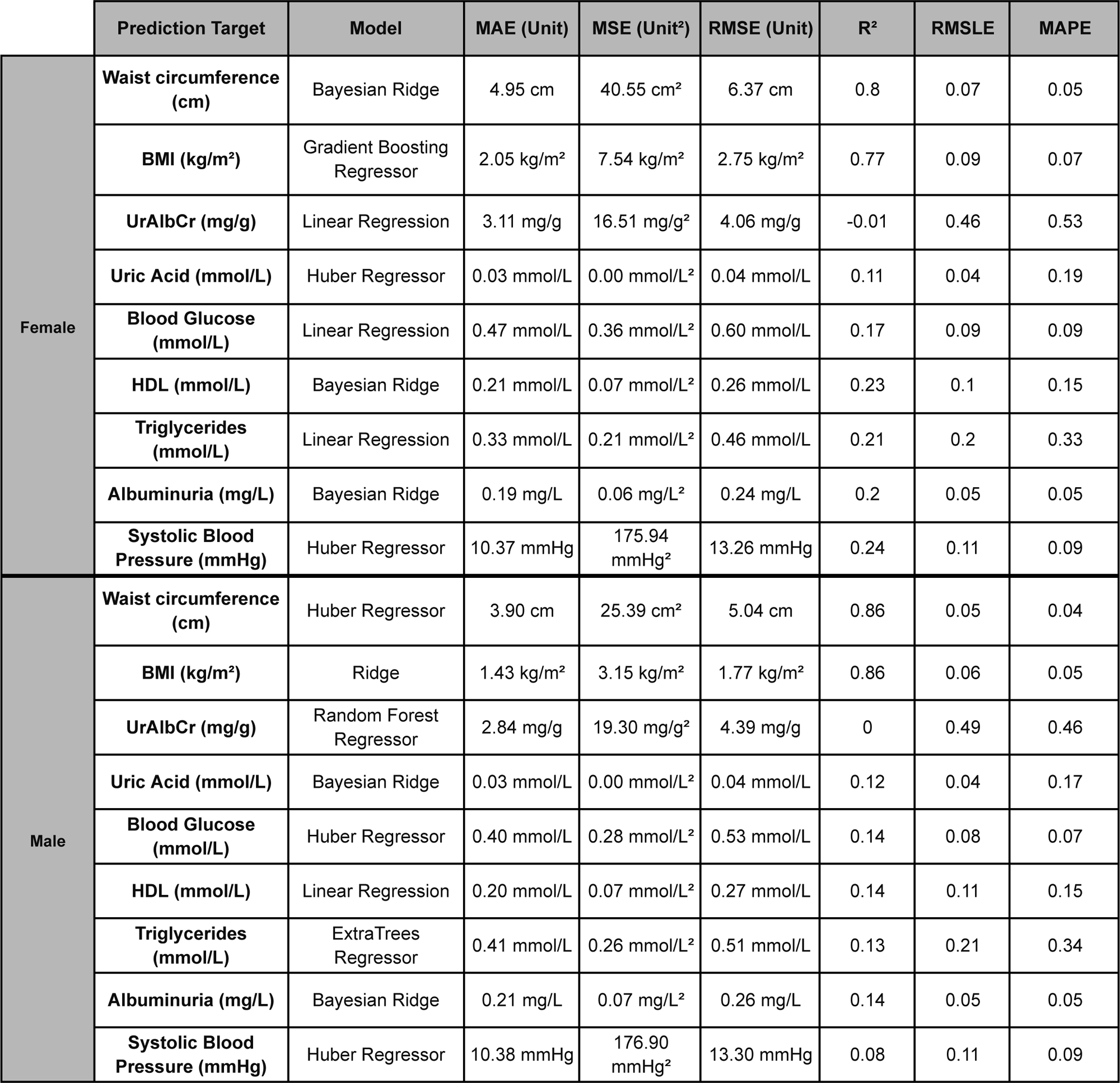
Best performing models for the various biomarkers in both male and female subgroups.

Table 6. Showcases the models devised for each biomarker, where the predictive outcome for each target was determined by the most optimal features alongside achieving optimal performance.

When examining Table 6, the top performing models for the female subgroup were waist circumference, BMI, blood pressure, blood glucose and albuminuria with the following evidence;

**Waist Circumference:** The Bayesian Ridge model recorded a Mean Absolute Error (MAE) of 4.95 cm and a Root Mean Squared Error (RMSE) of 6.37 cm. The Mean Squared Error (MSE) was relatively high at 40.55 cm². The R-squared (R²) value was 0.8, and the Root Mean Squared Logarithmic Error (RMSLE) and Mean Absolute Percentage Error (MAPE) were 0.07 and 0.05, respectively. The validation metrics when combined suggested that while the model generally performed well in estimating waist circumference (cm), there remains room for improvement in predicting actual values that fall significantly outside of observed mean range 93.47 (±14.20) cm as indicated by the elevated MSE.

**BMI:** The Gradient Boosting Regressor model showed an MAE of 2.05 kg/m², RMSE of 2.75 kg/m², and an MSE of 7.54 kg/m². The R-squared value was 0.77, and the MAPE was 0.07. Overall, these metrics indicated that the Gradient Boosting Regressor model performed reasonably well in predicting the target variable, with a relatively low error rate and a good level of explained variance.

**Blood Glucose:** The Linear Regression model for blood glucose had an MAE of 0.47 mmol/L, RMSE of 0.60 mmol/L, and an MSE of 0.36 mmol/L. The R-squared value was 0.17. Overall, these metrics suggested that while the Linear Regression model provided predictions for blood glucose levels, its accuracy and ability to explain the variance in blood glucose levels were relatively modest.

**Albuminuria:** The Bayesian Ridge model for albuminuria showed an MAE of 0.19 mmol/L, RMSE of 0.24 mmol/L, and an MSE of 0.06 mmol/L. The R-squared value was 0.20, and a MAPE of 0.05. Overall, these metrics suggested that the Bayesian Ridge model for albuminuria provided reasonably accurate predictions, capturing a notable portion of the variability in the data.

**Systolic Blood Pressure:** The Huber Regressor model for systolic blood pressure showed an MAE of 10.37 mmHg, RMSE of 13.26 mmHg, and an MSE of 175.94 mmHg. The R-squared value was 0.24 and a MAPE of 0.09. On average, its predictions deviated by about 10.37 mmHg, with a weaker fit to the data.

The male subgroup had the same top performing biomarkers as the female subgroup. Both the models for waist circumference and BMI indicated that they provided accurate predictions with a strong fit to the data, explaining a significant portion of the variability in waist circumference and BMI.The remaining top performing biomarkers were as follows;

**Blood glucose:** The Huber Regressor model showed an MAE of 0.40 mmol/L, RMSE of 0.53 mmol/L, MSE of 0.28 mmol/L. The R-squared value was 0.14 and MAPE of 0.07. Overall, these metrics indicated that while the Huber Regressor model provides acceptable predictions for blood glucose, its performance has room for improvement. It explained only a small portion of the variability in the target variable, and its predictions had a noticeable average deviation from the actual values.

**Albuminuria:** The Bayesian Ridge model’s performance in predicting albuminuria was evaluated with an MAE of 0.21 mmol/L, RMSE of 0.26 mmol/L, and MSE of 0.07 mmol/L, resulting in a R-squared value of 0.14 and a MAPE of 0.05. These findings suggested that the model demonstrates strong predictive ability for the target variable with minimal errors and high accuracy overall.

**Systolic blood pressure:** The Huber Regressor model for systolic blood pressure yielded an MAE of 10.38 mmHg, RMSE of 13.30 mmHg, and MSE of 176.90 mmHg. The R-squared value was 0.08 and the MAPE was 0.09. These results indicated that the model was well-suited to predict values within the first, second, and third quartile ranges but faces challenges in accurately predicting values outside of these ranges.

### 3.3 Validation test of actual results vs predictive results

After creating and refining the prediction models, a validation test was performed on the hold out set (Test Data) for both male (n=119) and female (n=121) groups for each of the biomarkers. In order to evaluate the predictive ability on the test set, we grouped the results within a 5% and 10% error respectfully. Heatmaps showing these results are displayed in Figures 4 and 5 for the female and male subgroups. Higher values (shown in yellow) demonstrated effective predictions by the optimised ML model, while blue indicated ineffective predictions.

**Figure 4:**
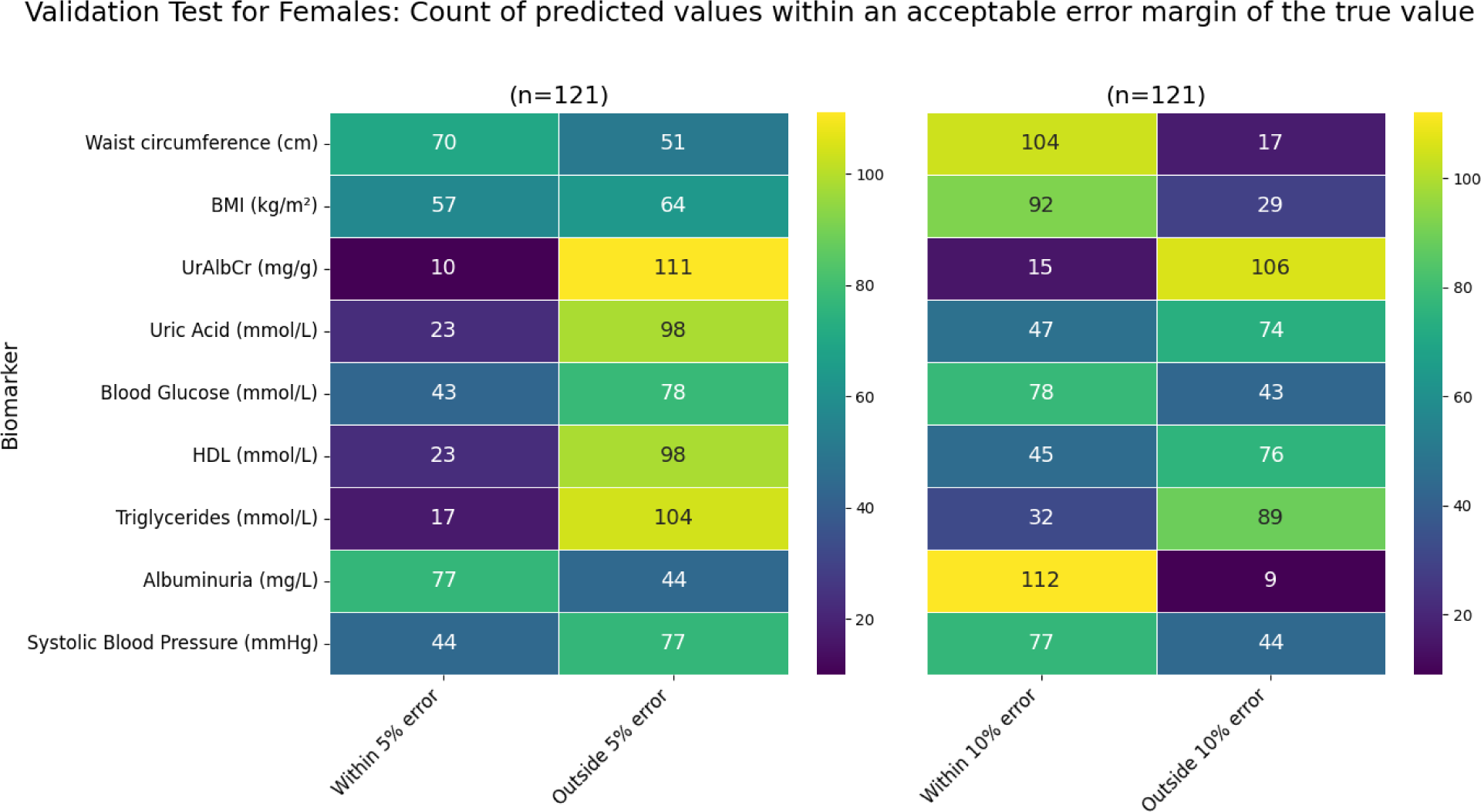
Heatmap showing the number of individuals where the predicted value falls either within or outside of a 5% and 10% error of the actual value in the female subgroup (from the n=121 test group).

**Figure 5:**
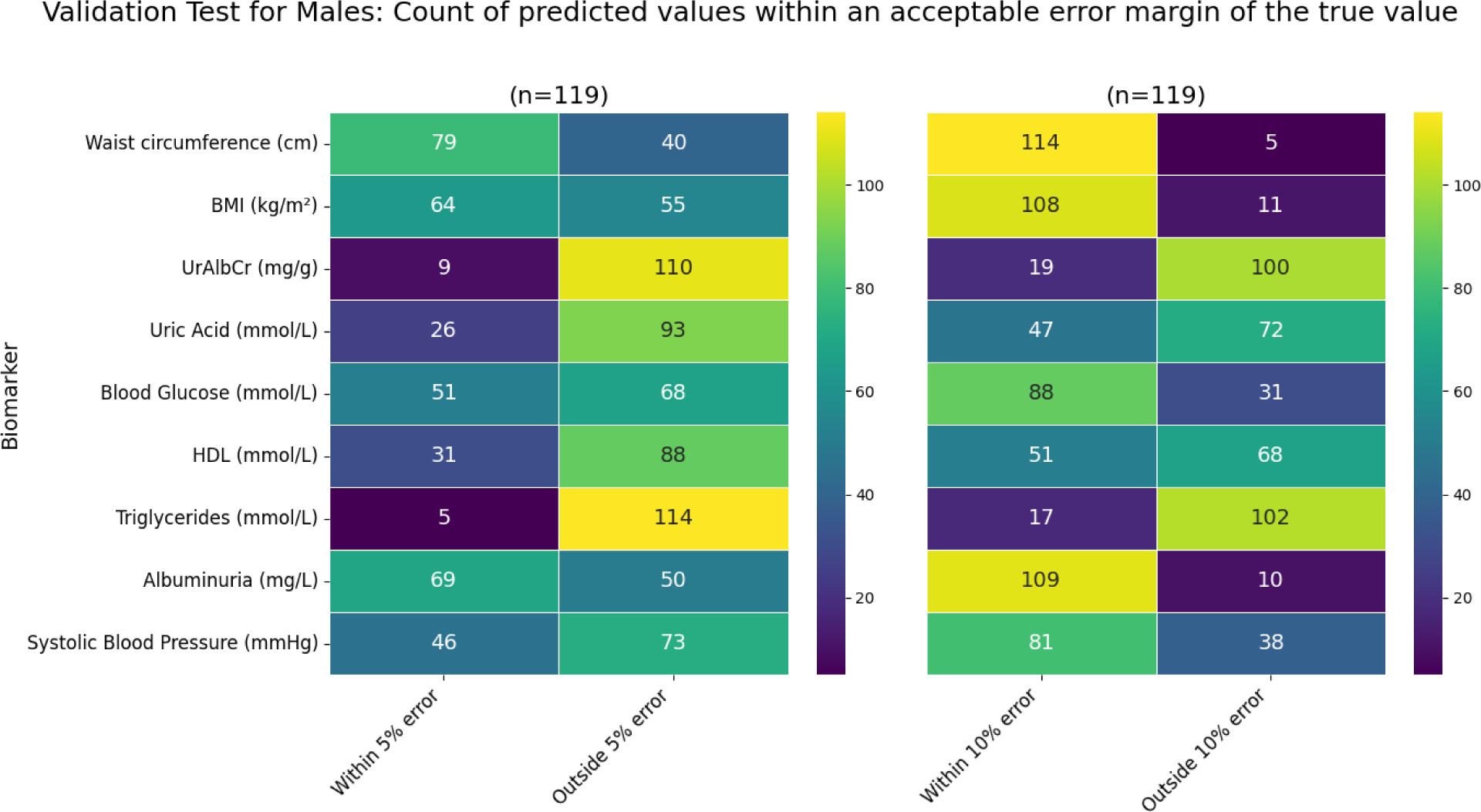
Heatmap showing the number of individuals where the predicted value falls either within or outside of a 5% and 10% error of the actual value in the male subgroup (from the n=119 test group).

As seen in Figure 4, the biomarkers that had the majority of individuals falling within the 10% error category were albuminuria, waist circumference, BMI, blood glucose, and systolic blood pressure. Of these biomarkers, the one consisting of the highest number of individuals was albuminuria with 93%, waist circumference with 86%, BMI with 76%, and the lowest two being blood glucose and systolic blood pressure with 64%. From a validation metrics point of view these were also the top performers with the exception of systolic blood pressure having a very high MSE value. The biomarker with the least number of individuals in the “within 10% error” category was UrAlbCr with 12%, followed by triglycerides (26%), HDL (37%), and uric acid (39%) respectively.

Despite a 10% error being acceptable for prediction purposes, in a clinical sense, that error could result in an individual falling incorrectly into an abnormal diagnostic range. For this reason a error of 5% was also evaluated and it can be seen that albuminuria resulted in 64% of the individuals falling within this range, followed by waist circumference (58%). UrAlbCr (8%) and triglycerides (14%) were the lowest two biomarkers respectively. Both uric acid and HDL had the same percentage (19%) of individuals for this category.

Upon examining Figure 5, the male subgroup had a higher percentage of individuals falling within the “within the 10% error” category compared to the female group overall; albuminuria (92%), waist circumference (96%), BMI (91%), blood glucose (74%), and systolic blood pressure (68%). The triglycerides had the lowest number of individuals (14%) followed by UrAlbCr (16%), and then uric acid (39%) and HDL (43%).

Within the 5% margin of error for the male subgroup, waist circumference exhibited the greatest number of individuals, followed by albuminuria. A similar pattern was noted in the female subgroup with these markers occurring in reverse order. BMI also had more than 50% of the individuals falling within this category which was different from the female group. Again, the lowest two biomarkers for this category were triglycerides and UrAlbCr, which was reversed in the female subgroup.

The results seen in the validation test were consistent with the validation metrics determined in Table 6.

## 4. Discussion

Our results indicated that in spite of the features being reduced as part of the RFECV method, the predictive ability of the ML models remained optimal in their ability to predict values within the mean range. The four biomarkers that need significant improvement in terms of model hyperparameters and feature selection for predicting biomarkers are triglycerides, HDL, uric acid, and UrAlbCr in both male and female subgroups.

### Lower performing markers

**Uric acid:** The methodology of this work was compared to the work done by Sampa et al. who were able to successfully predict uric acid using a similar approach to this study. In this study a RMSE of 0.04 mmol/L for both subgroups was obtained and was similar with the findings reported by Sampa et al. who had a RMSE of 0.03 mmol/L, using a boosted decision tree model. A notable distinction between this study and that of Sampa et al. pertained to the input features employed for uric acid prediction. Sampa et al. utilised a set of 17 features, comprising 12 core clinical measurements, 3 socio demographic attributes, and 2 dietary parameters. Of these, five features overlapped with those employed in the female subgroup of our study, namely sex, BMI, age, waist circumference, and blood glucose. In the male subgroup, only BMI coincided with the features employed by Sampa et al. [21]. The models they used were boosted decision tree regression, decision forest regression, bayesian linear regression, and linear regression. They found the boosted decision tree regression to be the best performer and are also considering adding work stress, daily physical activity, alcohol intake, and eating red meat as additional features for improving prediction.Their cohort was a Bangladesh cohort and their data was not stratified according to biological sex. In this study it was found that huber regressor in females and bayesian ridge in males were the favoured options based on the machine selection. The three differences between the two studies were the feature selections, the best machine model as well as the split between the male and female cohort.

**Triglycerides and HDL:** Triglycerides and HDL did not perform optimally in this study, the research done by Palumbo et. al had contradictory favourable prediction results for triglycerides, total cholesterol, HDL and urea while having a lower accuracy for glucose and total proteins. In their methodology they used 100 k-folds of training to improve model efficiency, whereas in this study 10 k-folds was used. Their process took 23 days using two 3.6 GHz Intel ® Core™ i7–4790 CPU systems running in parallel. In terms of the ML models, like this study, Palumbo et al also used regression models such as partial Least Squares Regression, and Support Vector Regression, Decision Tree, and Random Forest with the exception of the unsupervised ML Neural Networks model and the ensemble method [22].

**UrAlbCr:** The final marker that did not perform optimally was UrAlbCr. The research done by Huang et.al had these features included in their model training; sex, age, BMI, duration of diabetes, smoking, alcohol, fasting plasma glucose, glycated haemoglobin, triglycerides, high-density lipoprotein cholesterol, low-density lipoprotein cholesterol, alanine aminotransferase, creatinine, systolic blood pressure, diastolic blood pressure. They found that creatinine level was the most important feature, followed by systolic and diastolic blood pressure, glycated haemoglobin, and fasting plasma glucose in predicting UrAlbCr. This study had similar features selected in predicting UrAlbCr. The markers that Huang et al selected differently were glycated haemoglobin, low-density lipoprotein, alanine aminotransferase and diastolic blood pressure. Similarly the research done by Huang et al. also trained using 10 k-folds, using ML methods, namely classification and Regression Tree, Random Forest, Stochastic Gradient Boosting, and Extreme Gradient Boosting.

### Higher performing markers

**Waist circumference:** The waist circumference model for females was able to predict 86% of the test cohort within a 10% range of the mean using the Bayesian Ridge model as observed. The MSE was relatively high at 40.55 cm^2^. The R-squared value was 0.8, indicating the model’s ability to explain around 80% of the variability in waist circumference. Moreover, the male subgroup exhibited comparable outcomes (predicted 96% within a 10% range of the mean) using the Huber Regressor model, with the exception of a significantly lower MSE value of 25.39 cm². Based on the results in this study it was hypothesised that sex-related differences significantly influence the importance of features in predicting waist circumference estimates using supervised ML algorithms. Apart from the sex-related differences, another factor which may be contributing to the differences between the two groups could be the diversity in race found in the NHANES dataset (Table 3). Zou and colleagues conducted a similar study. Their cohort was a mixed population of Asian, Black, White, Hispanic and Chinese. The models they used were Extreme Gradient Boost, Semi-Bayesian Ridge Regression, Bozeman Linear Regression and Linear Regression and they assessed these models by splitting the data according to combined cohort, males overall and split according to race, and females overall as well as splitting it according to race. They used a 10-fold training with their data split 90% training and 10% test. Their input variables included height, weight, BMI, age, race and sex, with BMI, weight and Asian features being the stronger contributors for waist circumference prediction. In their results they found the Extreme Gradient Boost model as the top performer for most of the categories they examined with the Linear Regression having a 0.01 RMSE difference for the overall male subgroup. Their sex-combined model performance (RMSE 4.70cm) did not differ significantly from the sex stratified models (Female: RMSE 5.41cm; Male: RMSE 4.05cm) which is contradictory to the results found in this study (Female: RMSE 6.37cm; Male: RMSE 5.04cm). Zhou and colleagues argued that combining the sexes was the way forward in future models for waist circumference. They also found there was a tendency to overestimate waist circumference in United Kingdom females which did not occur in the United States females and suggested that a sociocultural factor could account for the difference between the two [23]. It was also found in our study that race contributed towards a poorer performance of the models, which was why it was removed as an input feature in the model training.

**BMI:** The BMI model in the female group was able to predict 76% within the mean range using the Gradient Boosting Regressor yielding validation metrics of MAE of 2.05 kg/m² and MSE of 7.54 kg/m², with a MAPE of 0.07. Conversely, for males, favourable results were achieved using the Ridge model with a 91% prediction and validation metrics showing a lower MAE at 1.43 kg/m² and MSE at 3.15 kg/m², along with a MAPE of 0.05, however, when comparing both sexes’ waist circumference in relation to BMI, it was evident that females had noticeably higher MSE values. The mean BMI for the female group was also noticeably higher 27.87 (±5.85) compared to males 27.39 (±4.82) contributing towards the larger MSE in females. Another difference noticed between the two subgroups was differences in the features selected for BMI. The features used to predict BMI in females included waist circumference, UrAlbCr, age, triglycerides, albuminuria, uric acid, blood glucose, systolic blood pressure, HDL, and smoking status. These selected features were significantly more comprehensive than the males which only included waist circumference and age. The results from the Levene’s test could possibly explain a reason for differences found between the features selected for each subgroup. Levene’s test revealed that while the average BMI might be similar between sexes, there are notable variations in BMI measurements between each group. Studies have shown that variance in data influences the feature selection and overall model prediction capability, therefore should be carefully considered in future work [24].

Delnevo et al. also did a study on predicting BMI with ML, however, they considered negative and positive psychological variables as their input features and after the BMI prediction they then applied it to an obesity classification model. Only focusing on the BMI prediction aspect of their study, the features they considered included depression, trait anxiety, binge eating, expressive suppression, happiness, emotional intelligence, and emotion regulation. They also had an 80% training and 20% test split for their data with a 4-fold training as their data set was too small for the usual 10-fold strategy. The ML models used were K-Nearest Neighbour, Classification and Regression Tree, Support Vector Machine, Multi-Layer Perceptron, Ada Boosting with Decision Tree, Gradient Boosting, Random Forest, Extra Tree, Lasso, and Elastic Net Regression. They found that their regression results between the positive and negative features were favourable with Gradient Boost. The negative features had an MAE of 4.34 which was almost 2-4 times higher than the results obtained in this study. This is indicative of a more favourable result obtained in this study [25]. Sancar and Tabrizi also conducted a study in estimating BMI using an Adaptive Neuro Fuzzy Inference System (neural network). Their cohort was obtained from the Near East University Hospital in Cyprus and also split according to biological sex with a 70%: 30% training and test data split. The features they considered included BMI, waist circumference, systolic blood pressure, diastolic blood pressure, fasting glucose, total cholesterol level, high-density lipoprotein, low density lipoprotein, triglycerides, uric acid, high sensitivity C-reactive protein, age, and homeostatic model assessment-insulin resistance. Their final model only included waist circumference, fasting glucose, high-density lipoprotein, triglycerides, and homeostatic model assessment-insulin resistance as the training features for both the male and female models. They obtained close results for the male and female observed BMI to predicted BMI values. The female RMSE results were 1.914 kg/m² and female BMI prediction results included 30.90 kg/m² (observed): 30.50 kg/m² (predicted), and 40.60 kg/m² (observed): 41.10 kg/m² (predicted), with the males having a RMSE result of 1.817 kg/m² and BMI prediction results of 37.20 kg/m² (observed): 37 kg/m² (predicted), and 31.50 kg/m² (observed); 31.80 kg/m² (predicted) [26]. The RMSE values obtained in this study for the females were 2.75 kg/m² being higher than Sancar and Tabrizi and the males had an RMSE value of 1.77 kg/m² which was lower than that of Sancar and Tabrizi, indicating that the female subgroup in this study had a lower prediction value whereas the males in this study had a higher prediction value compared to the Sancar and Tabrizi study. Compared with these two studies, the BMI biomarker performed relatively well with the male BMI prediction performing the best.

**Blood glucose:** In the female subgroup, the Linear Regression model was selected as the most effective predictor for blood glucose, demonstrating an MAE of 0.47 mmol/L, RMSE of 0.60 mmol/L, MSE of 0.36 mmol/L, and MAPE of 0.09. It had a prediction value of only 64% within the 10% range of from the mean for the test group. Additionally, the optimal features selected for the blood glucose model identified four key features: age, systolic blood pressure, waist circumference, and HDL. In the male subgroup, the Huber Regressor model produced an MAE of 0.40 mmol/L, RMSE of 0.53 mmol/L, and MSE of 0.28 mmol/L with an R-squared value of 0.14 and a MAPE of 0.07. The features selected included age, BMI, UrAlbCr, and triglycerides. These measurements indicated that the Huber Regressor model was able to deliver sufficient prediction results with these features for blood glucose with a 74% predictability for the test group within the 10% range of the mean. Although it performed better than the female group, there is room for improvement in overall performance especially in the medical context. In comparison to literature, Fu et. al did a classification approach to blood glucose prediction in terms of glycaemic control. The regression models they applied were K-Nearest Neighbours, Logistic Regression, Random Forest, Support Vector Machine, and Extreme Gradient Boost. They found extreme Gradient Boost to be the preferred model for predicting blood glucose concentrations. The features they considered for training included age, sex, experimental grouping, family history, education level, dietary assessment, complications (retinopathy, kidney disease, peripheral neuropathy, peripheral atherosclerosis, intermittent claudication), hypertension, drinking status, smoking status, BMI, pulse rate, and key biochemical indicators (HDL, Hb, K, Na, Cl, CO2, Ca, P, AKP, GPT, GOT, rGT), however, they did not stipulate the final features selected for training [27]. Van Doorn et al’s study considered predicting blood glucose at 15 minutes and 60 minutes intervals. The ML models they used included Autoregressive Integrated Moving Average, Support Vector Regression, Gradient Boosting Systems, Shallow and Deep Multi-Layer Perceptron, Neural Networks, Recurrent Neural Network, and Long Short-Term Memory Networks. The models were trained on continuous glucose monitoring data. The Recurrent Neural Network predicted glucose accurately for the 15 minute window and the Long Short-Term Memory Network was able to predict glucose accurately for the 60 minute time frames for all the participants in the normal glucose, prediabetes and diabetes ranges. The RMSE values they obtained were 0.477 mmol/L and 0.922 mmol/L respectively [28]. The RMSE values obtained in this study were comparable with the Van Doorn study with the females having an RMSE of 0.6 mmol/L and males with an RMSE of 0.53 mmol/L.

**Albuminuria:** The albuminuria biomarker in the female subgroup, used the Bayesian Ridge model as the model of choice and was able to predict 93% of the test subjects within the 10% mean range. The validation metrics obtained were a MAE of 0.19, RMSE of 0.24 mg/L, and a R-squared value of 0.2. The model appeared to effectively capture approximately 20% of the variance in the albuminuria levels. With low RMSLE and MAPE values of 0.05 each, the model demonstrated consistent predictions. In the male subgroup albuminuria demonstrated an optimal fit also using the bayesian ridge model with an MAE of 0.21 mg/L, MSE of 0.07 mg/L, and RMSE of 0.26 mg/L signifying accurate predicted values as indicated by a minimal error confirmed by MAPE at 0.05 and the prediction of the test group with a 92% prediction within the 10% mean range. Khitan et. al conducted a study that utilised a ML classification method to predict albuminuria. They took into account several features such as age, HDL, triglycerides, and creatinine, all of which were also employed in this research. The Random Forest Classifier, with a true positive score of 87% emerged as the most effective model for the classification technique employed [29].

**Systolic blood pressure:** This study revealed the Huber Regressor to be the most effective model for predicting systolic blood pressure in females, yielding a MAE of 10.37 mmHg, a MSE of 175.94 mmHg^2^, and a RMSE of 13.26 mmHg. The model predicted values of 64% within a 10% range to the mean female patients. In the male group, the Huber Regressor was also identified as the most suitable model for the dataset, achieving a MAE of 10.38 mmHg, MSE of 176.90 mmHg², and RMSE of 13.30 mmHg, suggesting its capability to predict values close to the actual mean value. In the test group it was able to predict 68% of the test group within 10% of the mean range. Including patients with hypertension may have introduced significant variability in the data resulting in a substantial deviation beyond the normal range indicated by the high MSE. This research also involved individuals with normal systolic blood pressure and stage 1 hypertension (130 to 139 mmHg), revealing that the average systolic blood pressure for women was slightly lower than that of men, at 116.59 (±15.15) mmHg compared to 121.84 (±13.49) mmHg for males. The impact of age on blood pressure is well acknowledged, with ageing leading to reduced flexibility in blood vessels, potentially resulting in higher blood pressure and increased risk of hypertension development, especially isolated systolic hypertension among those aged 50 years or older [30]. Age played a prominent role as an input feature across most models developed in this study (Figure 3), however, it may have also contributed to the observed error as models were only aggregated by sex and not by age. A hypothesis is that stratifying the data according to age and hypertension range for both subgroups would improve the prediction capability of this biomarker. A study conducted by Zheng and Yu used a variety of different features in their model training.These features included; sex, marital status, smoking status, age, weight, overweight, height, BMI, exercise level, alcohol consumption, stress level, and salt intake level. The models they considered were Linear Regression, Support Vector Machine, Decision Tree Regression, Gaussian Process Regression, and Artificial Neural Network. The models in this study performed equally well compared to the Neural Network utilised in Zheng and Yu’s study, however, still require optimising in order to be accepted accordinging to ANSI/AAMI standards which are between 5 and 8 mmHg [31].

**Limitation:** A limitation observed in this study was the impact of age on the data. This was particularly evident in the case of the BMI biomarker, which showed a higher prevalence of elevated BMI in the 20-29 age range compared to the 60-69 age range. To enhance the predictive capabilities of the ML model, one approach could be to stratify the data based on different age ranges which are diagnostically relevant. Another hypothesis is that this study did not stratify biomarkers according to diagnostic ranges, such as normal vs hypertensive readings for blood pressure. Training our model using specific diagnostic ranges for each biomarker may lead to more accurate predictions within those particular health conditions. The conventional approach to ML methodology typically involves the pre-selection of specific models. However, this study departs from this by adopting a more thorough approach by selecting multiple models without bias and facilitating their competition for optimal performance confirmed using the validation metrics. It was also noticed that features contributed towards the accuracy of a model’s prediction capability and therefore special consideration towards feature selection is a vital part of ML hyperparameter tuning.

**Future works:** Although a wide variety of validation metrics were used to analyse the various biomarkers, future works would include selecting a main metric with a supporting metric that suits the requirements for each biomarker. For example, for models where outliers are important in medical ranges such as blood pressure, squared metrics should feature e.g RMSE. For models with expected linear relationships such as BMI or waist circumference, metrics such as R^2^ or MAE would be a better fit. A surprising observation is that linear regression models were primarily selected by ML as optimal models (Table 5). As shown in the above examples, non linear regression models are a typical choice such as Random Forest. These findings could be due to the small data set. To obtain a more accurate training set a larger (a database of at least 10000) data size and a wider variance would need to be sourced in future work. As outlined in the methodology, the ML model was initially trained to distinguish between male and female data. Future investigations will explore the impact of combining these groups and compare the biomarker prediction capabilities with the results obtained in this study. Additionally, two other comparisons will be made: one to evaluate the effect of stratifying the data by age groups on biomarker prediction, and another to assess the impact of categorising the data according to diagnostic ranges or groups. The motivation for this was inspired by the results found in the Levene, t-test and feature selection results, where it was clearly seen how data variability between the two subgroups contributed towards feature selection and therefore influenced the model prediction capabilities for the various biomarkers. Another aspect to consider in biomarker prediction optimisation would be to include other features shown to contribute to triglyceride, HDL, uric acid, and UrAlbCr prediction improvement. These markers include glycated haemoglobin, c-reactive protein, presence of diabetes, hypertension, diastolic blood pressure. Once the prediction models of these have been improved, the next step would be to use the prediction capabilities of these models in inferring markers with a low error as well as infer multiple markers from one feature. This would be valuable in giving humans insight into valuable biological markers when limited data is available.

## 5. Conclusion

In conclusion, this study utilised a supervised ML framework to predict waist circumference, BMI, albuminuria, UrAlbCr, uric acid, blood glucose, HDL, triglycerides, and systolic blood pressure (mmHg) based on 12 input variables. The RFECV method facilitated the optimal selection of features in the prediction model optimization process. Following model training, Bayesian Ridge, Ridge, and Linear Regression emerged as the most frequently employed models. Notably, biomarker models for BMI, waist circumference, blood glucose, systolic blood pressure, and albuminuria demonstrated acceptable performance with poorer predictions found with triglycerides, HDL, uric acid, UrAlbCr for both male and female subgroups. Overall the male subgroup had higher prediction scores than the female subgroup. This study showed the ability of ML models to predict complex biological markers and the combination of features towards said predictive models being impacted by sex. It also highlighted the critical importance of selecting appropriate features and properly stratifying data (such as according to sex), as these factors significantly influence marker prediction capability.

## 6. Acknowledgements

The authors would like to acknowledge Renier van Rooyen for his contribution and feedback in examining the ML methodology and results in the article.

